# ASCENT: Annotation-free Self-supervised Contrastive Embeddings for 3D Neuron Tracking in Fluorescence Microscopy

**DOI:** 10.1101/2025.07.23.666425

**Authors:** Haejun Han, Robert Pritchard, Hang Lu

## Abstract

Tracking individual neurons in dense 3D microscopy recordings of fast-moving, deforming organisms is a critical challenge in neuroscience. This task is often hindered by complex tissue motion and the need for laborious manual annotation. Here, we introduce ASCENT, an annotation-free computational framework that learns robust representations of neuronal identity that are invariant to tissue deformation. ASCENT trains a Neuron Embedding Transformer (NETr) through a self-supervised contrastive scheme on augmented data that mimics tissue deformations. NETr generates highly discriminative embeddings for each neuron by combining visual appearance with positional information, refined by the context of all other neurons in the frame. On a challenging benchmark *C. elegans* datasets, ASCENT achieves state-of-the-art tracking accuracy, surpassing supervised methods. The method is robust to image noise, highly data-efficient, and generalizes across different imaging conditions. By eliminating the annotation bottleneck, ASCENT provides a practical and scalable solution for the robust analysis of whole-brain neural dynamics in behaving animals.

## Introduction

Understanding how the activity of large neural populations gives rise to complex behaviors is one of the central goals in neuroscience. Functional imaging, particularly in behaving and moving animals, provides a way to directly associate the dynamics of activities of neurons and circuits to relevant behaviors. Advances in volumetric microscopy, such as light-sheet or light field microscopy and sculpted light techniques, now allow one to record three-dimensional (3D) time-lapse images of nearly entire brains in small model organisms such as *C. elegans* [1–3] and larval zebrafish [4–7] at single-cell resolution. Extracting biological insight from these rich datasets, however, first requires solving the critical task of multi-neuron tracking: accurately following the position and identity of each neuron through time to extract its individual activity trace [8–11].

This tracking problem is exceptionally challenging in the context of 3D fluorescence microscopy. Neurons are often densely packed and, when viewed in isolation, lack uniquely distinguishing visual features like texture or shape, making them appear visually homogeneous [9, 10]. They are also subject to complex, non-rigid deformations as the animal moves and its brain bends and stretches [12–14]. To address this, early computational methods relied on registering point clouds representing neuron coordinates between frames [9, 15] or aligning them to a reference atlas [16–18]. Subsequent deep learning techniques often focused primarily on positional features [19, 20], sometimes overlooking valuable image content from the local cellular neighborhood that can provide context for disambiguation. More recent methods incorporating image features have improved accuracy by using techniques such as graph matching [21], voxel classification [22], or local and global image registration [13, 23]. However, despite these advances, many of these state-of-the-art methods still do not yield high enough accuracy to avoid extensive manual corrections for tracking, in part because of noises in fluorescence imaging, imperfections in object detection, and large motions. Additionally, creating the ground-truth data for supervised learning (i.e. manually identifying and linking hundreds of neurons across thousands of volumetric frames) is an extremely labor-intensive and expertise-dependent process that limits the scalability and widespread adoption of these powerful methods [24–26].

Here, we present Annotation-free Self-supervised Contrastive Embeddings for Neuron Tracking (ASCENT), a computational framework designed to overcome these challenges by learning embeddings directly from unlabeled imaging data. We engineer a Transformer-based network that synergistically integrates local visual appearance with positional information to track dense, visually similar cells. This network uses self-attention to refine each embedding with contextual cues from all co-detected neurons, providing robustness against the complex, non-rigid deformations inherent to imaging behaving animals. Importantly, we train this network using a self-supervised contrastive learning scheme with a tailored data augmentation strategy, which completely eliminates the need for manual track annotations. We demonstrate that the synergistic use of both visual and positional features is critical for robust performance, particularly in noisy conditions, which is a common feature in 3D video microscopy for moving/deforming samples. Our demonstrations on both a public benchmark and a newly acquired dataset of whole-brain imaging of spontaneous activities in freely moving *C. elegans* and odor-stimulated *C. elegans* show that ASCENT surpasses state-of-the-art accuracy and generalizes across different imaging conditions. Furthermore, ASCENT is highly data-efficient, achieving high performance with as few as eight unlabeled training frames. Together, these features establish ASCENT as a practical and scalable framework that enables robust downstream analysis of neural dynamics in behaving animals in wide-ranging scenarios.

## Results

### The ASCENT framework for annotation-free neuron tracking

Tracking individual neurons in dense, deforming 3D volumes is a great challenge in fluorescence video microscopy, particularly in freely moving animals. Neurons often appear visually similar, and the sample is subject to large, complex movements that include non-rigid deformations, where different parts of the tissue can shift and warp independently (Figure 1a,b). To overcome this challenge, we developed ASCENT (Figure 1c-e), a computational framework that transforms the tracking problem into a feature-matching task [27]. The core principle of ASCENT is to train a deep embedding network that generates a unique and discriminative feature vector, or embedding, for each neuron at every timepoint. This transforms the complex task of spatiotemporal linking into a more straightforward feature-matching problem between different frames.

**Fig. 1:**
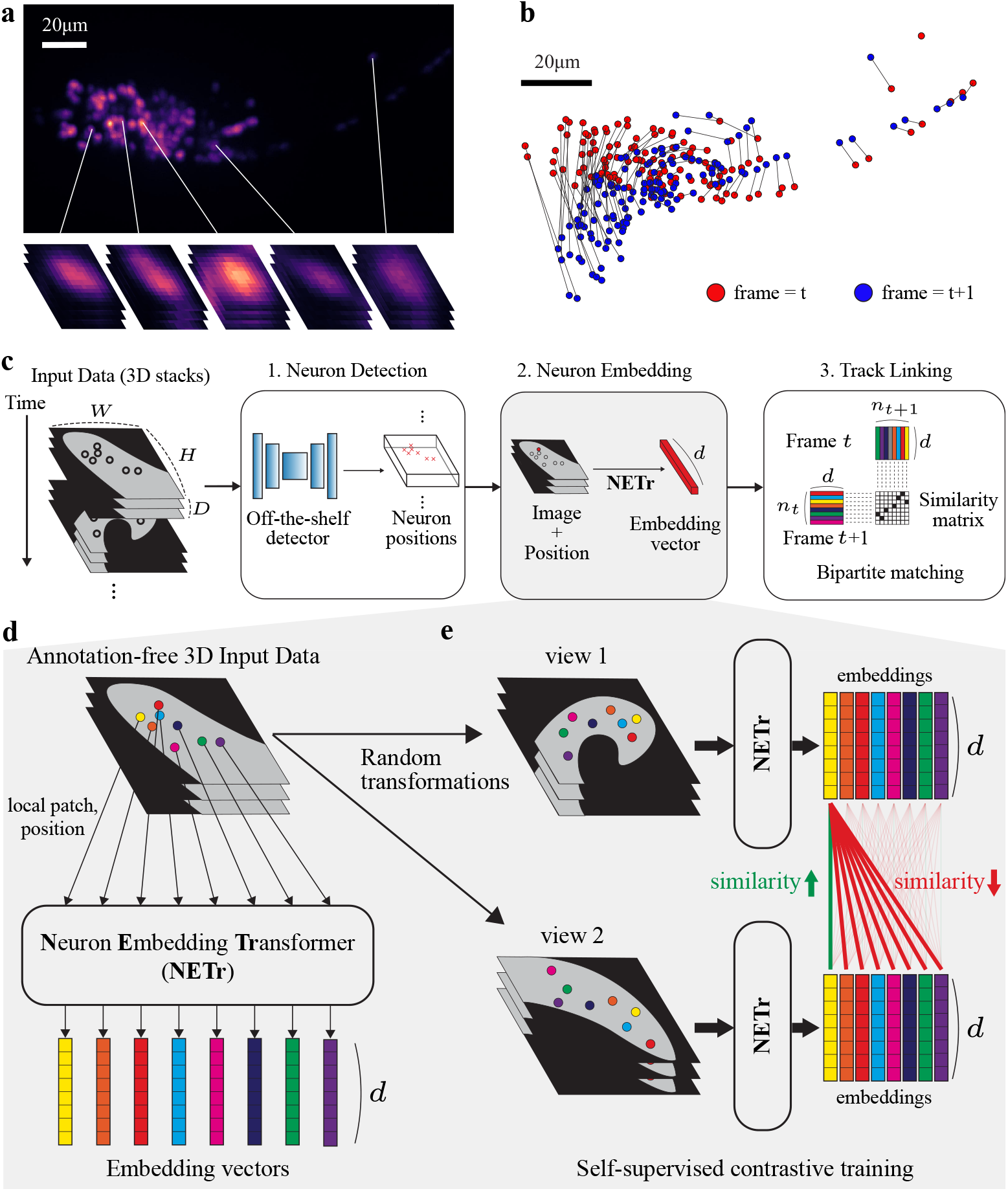
The ASCENT framework for annotation-free neuron tracking. **a,b**, Challenges in 3D neuron tracking, including visually indistinguishable neurons in a dense cluster (**a**) and complex, non-rigid frame-to-frame movements (**b**). **c**, The ASCENT pipeline proceeds from neuron detection in input 4D data to neuron embedding via NETr, and finally to track linking. **d**, Conceptual diagram of the Neuron Embedding Transformer (NETr) in the neuron embedding step. NETr takes a 3D image and detected neuron positions as input to produce a unique, d-dimensional embedding vector for each neuron. **e**, Conceptual diagram of the self-supervised training scheme. For a given frame, two different, randomly augmented “views” are generated and passed through the embedding network. A contrastive loss then ensures that the embeddings for the same neuron are consistent across views, while embeddings for different neurons are pushed apart.

The framework is a modular three-stage process (Figure 1c). In the first stage, initial neuron coordinates are provided by an off-the-shelf detection algorithm (see Methods for details). This design choice makes ASCENT agnostic to the specific detection method and allows it to readily incorporate future advances in object detection or segmentation. In the second and most critical stage, our Neuron Embedding Transformer (NETr) computes a robust, context-aware representation for each detected neuron. In the third and final stage, neurons are linked across frames using a simple bipartite matching algorithm. This straightforward approach is made possible by the highly discriminative nature of the learned embeddings and avoids the complex post-processing and case-by-case parameter tuning required by many other tracking methods.

The NETr architecture is specifically designed to produce context-aware embeddings by synergistically combining local visual appearance and the relative position of neurons (Figure 1d, Supplementary Figure 1a; see Methods for details). While individual neurons often appear visually homogeneous, we hypothesized that the local cellular neighborhood provides a rich, identity-preserving visual signature. By extracting features from a 3D image patch around each neuron, a powerful visual backbone can learn these unique contextual patterns. However, visual features alone are insufficient for robust tracking under all conditions; they are inherently sensitive to image noise and can fail when spatially distant neurons have visually indistinguishable local surroundings. To complement this, NETr separately computes positional features from the coordinates, providing a global spatial frame of reference that robustly disambiguates such cases. These two feature modalities are combined and then processed by a Transformer encoder. The choice of this architecture was inspired by the success of its self-attention mechanism in the feature-matching literature [27–30]. This mechanism refines each neuron’s embedding by integrating information from all other neurons present in the same frame.

To overcome the manual annotation bottleneck inherent to supervised neuron tracking, we adopted a self-supervised contrastive learning scheme (Figure 1e, Supplementary Figure 1b; see Methods for details) based on the SimCLR framework [31]. ASCENT’s contrastive learning scheme draws on the success of contrastive methods in other contexts [32–35]. The network learns by being shown two different, randomly augmented “views” of the same set of neurons from a single frame. A contrastive loss function then trains the network to produce consistent embeddings for the same neuron across the two augmented views, while simultaneously pushing apart the embeddings of different neurons. Through this process, NETr learns to generate feature embeddings that are robust to deformations and imaging variability, yet highly specific to individual neuron identity, which allows ASCENT to track with better accuracy, with no manual annotations.

### An experimental condition-aware data augmentation strategy creates a discriminative embedding space

To learn features that are invariant to imaging noise and deformation, yet specific to each neuron, we implemented a data augmentation strategy that creates diverse and challenging views of the same underlying neurons while preserving their core identity (Figure 2a, Supplementary Figure 2). For instance, transformations such as rotation and elastic deformation are meant to mimic moving samples during imaging, while position jitter and color (intensity) jitter mimic perturbations due to the nature of the reagents and noise from the microscopy system.

**Fig. 2:**
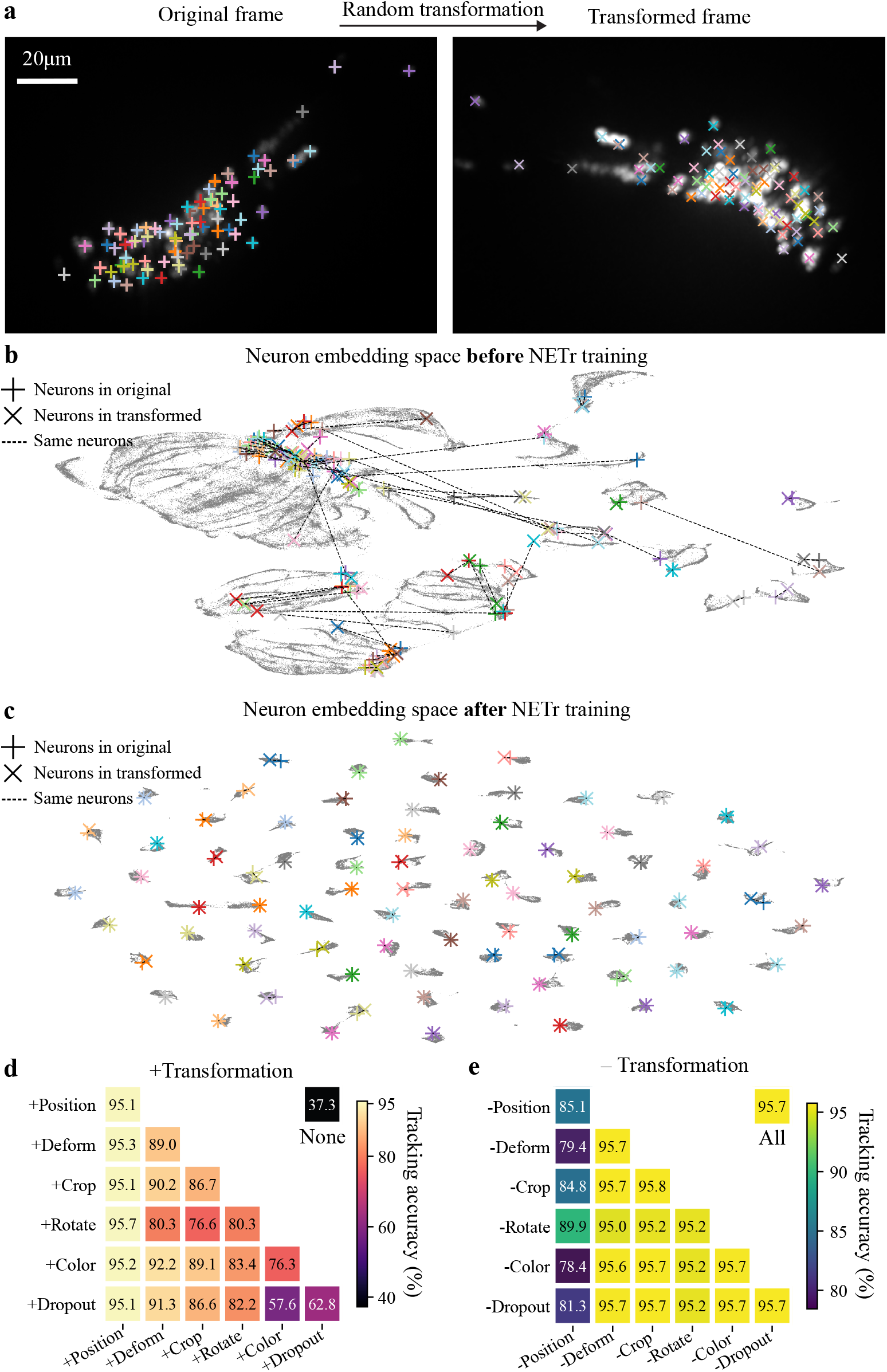
An experimental condition-aware data augmentation strategy creates a discriminative embedding space. **a**, Example of a random data augmentation applied to a frame from the NeRVE dataset. The original neuron positions are marked with ‘+’ and the transformed positions with ‘× ‘. **b**, UMAP visualization of the neuron embedding space before training. Colored markers represent neurons from the original (+) and transformed (×) views shown in **a**; gray points represent all other neurons. Dotted lines connect corresponding neuron pairs, which are scattered randomly across the space. **c**, Neuron embedding space after self-supervised training. The embeddings for corresponding neuron pairs are now tightly co-localized. **d,e**, Quantitative analysis of augmentation contributions on the NeRVE dataset. Heatmaps show the tracking accuracy gain when adding one or two transformations to a baseline (**d**), and the accuracy drop when removing one or two transformations from the full set (**e**), confirming the positive contribution of each component.

To understand how this process enables learning, we visualized the neuron embedding space before and after training for a whole-brain recording of a freely-moving *C. elegans* from a previous study [3] using uniform manifold approximation and projection (UMAP) [36] (see Methods for details). Before training, the embedding space is largely unstructured. While the pre-trained visual backbone provides some initial feature separation, the embeddings corresponding to most neuron identities are intermingled and indistinct (Supplementary Figure 3a). Consequently, the representations of the same neuron from an original and an augmented view are scattered randomly across the space (Figure 2b). This demonstrates that the untrained network is largely incapable of distinguishing between neurons, which would make any attempt at tracking via feature matching ineffective. After training with the contrastive objective, the embedding space becomes highly organized. Across the entire dataset, this results in the formation of tight, well-separated clusters, where each cluster is composed of the embeddings of a single neuron tracked through time (Supplementary Figure 3b). The embeddings for corresponding neuron pairs from the two views are pulled into close proximity, forming tight, identity-specific clusters that are well-separated from one another (Figure 2c). The emergence of this structure is a direct visualization of the learning process: it demonstrates that the network has successfully learned to generate a unique and robust feature signature for each neuron, creating a space where simple distance-based matching can be used for reliable tracking.

To determine the contribution of each transformation to this outcome, we systematically dissected the augmentation pipeline using the same dataset. We first trained the embedding network using only one or a combination of two transformations from our full suite of six to assess their individual and pairwise contributions (Figure 2d). While any single augmentation dramatically improved performance over the no-transformation baseline (37.3% accuracy), we found that applying random jitter to the neuron positions provided the single most effective gain, boosting tracking accuracy to 95.1%. This result is particularly significant; it demonstrates that the most critical factor in learning a useful embedding space is to force the network to be robust to small positional perturbations, which are inevitable in real data (variations from worm to worm, from frame to frame) and realistic detection pipelines. Furthermore, a combination of just two transformations – position jitter and random rotation – was sufficient to achieve the peak accuracy of 95.7%, matching the performance of the full suite of six augmentations.

To assess the relative importance and potential redundancy of each augmentation, we next performed the converse experiment, systematically removing one or two transformations from the full suite (Figure 2e). As expected, removing position jitter alone caused the most substantial performance drop, reducing accuracy from 95.7% to 85.1% (Figure 2e). In contrast, the removal of other augmentations (e.g. elastic deformation or color jitter) from the full set resulted in negligible changes to accuracy, suggesting a degree of functional overlap of these augmentations with each other (Figure 2e). This apparent redundancy likely arises because the inherent frame-to-frame variability in this challenging dataset provides a learning signal similar to that simulated by these specific transformations. It is worth noting that with other datasets, the contributions of these features may be quantitatively different and less redundant. Since no combination of augmentations meaningfully hampered performance, our results suggest that while a minimal set is highly effective for this dataset, the full suite could provide a robust default strategy that may confer additional benefits on more complex or varied datasets. Here, by transforming raw image data and neuron positions into a rich set of augmented views, the self-supervised training strategy successfully guides the network to learn an embedding space structured by neuron identity, which is required for accurate tracking.

### Synergistic integration of visual and positional features ensures tracking robustness

A common assumption in tracking visually similar cells is that local appearance is insufficient for reliable identification, which is why most methods primarily rely on positional information [9, 19, 20]. Our approach challenges this position-centric view, based on the hypothesis that the local visual context provides a rich source of identity information. While individual neurons may appear homogeneous, the 3D spatial arrangement of their neighbors within a local patch creates a unique geometric signature. We reasoned that even under non-rigid deformation, this local constellation of neurons provides a robust, identity-preserving feature that a powerful visual backbone can learn. However, visual context alone can be ambiguous under heavy image noise or when spatially distinct neurons share similar local neighborhoods. Thus, we designed the architecture of ASCENT to include both visual features and positional features. To dissect contributions of visual and positional features to ASCENT’s performance, we evaluated ablated models that used only visual features (‘Visual features only’) or only positional embeddings (‘Positional features only’) against the complete ‘Standard’ NETr model.

Visual features are highly susceptible to image noises, which are common and microscope-dependent. We thus, simulated various noise levels by adding zero-mean pixel-wise Gaussian noise with standard deviation *σ* applied to the normalized whole-brain recording of the freely moving *C. elegans* (Figure 3a,b; see Methods). On the original, low-noise imaging data, the standard model achieved the highest accuracy. Surprisingly, the model using exclusively local visual features performed nearly as well as the complete model. This demonstrates that our self-supervised learning strategy can extract powerfully discriminative information from local appearance alone (Figure 3b, red dots). However, this reliance on visual information is fragile; as noise levels increased, the accuracy of the visual-only model degraded severely. In contrast, the model using only positional information proved highly robust to this image degradation, maintaining its performance as image quality decreased (Figure 3b, blue dots). These results highlight the complementary nature of the two feature modalities. While visual information provides highly discriminative features under ideal conditions, positional information provides an essential scaffold of robustness against image noise. Therefore, the integration of both is necessary to achieve consistently high tracking accuracy across the variable conditions encountered in live-animal microscopy.

**Fig. 3:**
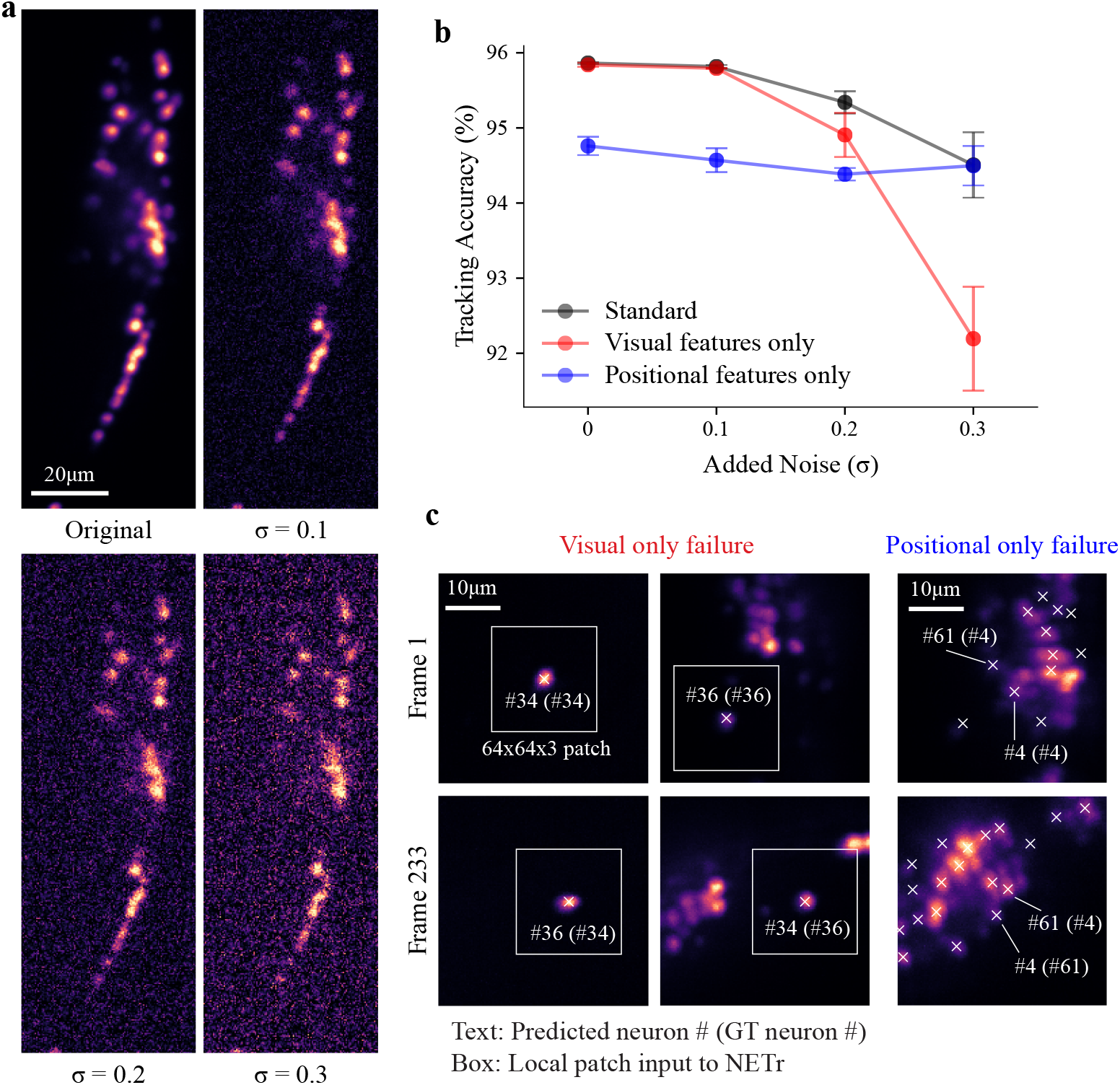
Synergistic use of visual and positional features enables robust tracking. **a**, Example of a frame from the NeRVE dataset with varying levels of synthetic Gaussian noise applied, from *σ* = 0.1 to *σ* = 0.3. **b**, Tracking accuracy of NETr architecture variants as a function of image noise. The ‘Standard’ model (black), using both feature types, achieves the highest accuracy in low-to-moderate noise conditions. The ‘Visual features only’ model (red) performs well on clean data but its accuracy degrades severely with noise. The ‘Position features only’ model (blue) is robust to noise even at extreme noise levels. Data are presented as mean *±* standard deviation over three independently trained models. **c**, Qualitative examples of the distinct failure modes of ablated models. Labels are formatted as “predicted neuron # (ground-truth neuron #)”. Left & Center: The ‘Visual features only’ model fails by swapping two visually similar neurons (#34 and #36) between frame 1 and frame 233. The white box indicates the 64×64 pixel local patch from which visual features are extracted. Right: The ‘Position features only’ model fails by swapping two spatially proximal but visually distinct neurons (#4 and #61) between frame 1 and frame 233.

Tracking errors visualized in Figure 3c further demonstrate the complementary nature of these components and their distinct failure modes: the visual-only model fails by swapping two visually similar but spatially separated neurons (#34 and #36) that are indistinguishable within their local image patch. In contrast, the position-only model fails by swapping two spatially proximal but visually distinct neurons (#4 and #61) whose relative positions become ambiguous in the presence of tissue deformation. This latter failure mode is a key limitation of trackers that largely ignore image content. By synergistically integrating both modalities, ASCENT’s full architecture overcomes the specific limitations of each, using positional encoding to disambiguate visually similar neurons and local appearance to resolve identities when relative positions are distorted.

### ASCENT sets a new state-of-the-art performance and generalizes across diverse datasets

We validated ASCENT’s performance on two datasets with different characteristics: a publicly available recording of a freely moving *C. elegans* (NeRVE) and an inhouse recording of a worm confined in a microfluidic channel subjected to chemical stimuli (Figure 4a). We first benchmarked ASCENT against state-of-the-art methods on the NeRVE dataset (Table 1, Figure 4b). ASCENT achieves a tracking accuracy of (95.90%), surpassing the previous best-performing supervised method, ZephIR (94.43%) [13] (Figure 4b). Very importantly, we note that this superior performance is achieved with zero manual annotation time, in contrast to the hours of expert annotation per video analyzed for supervised methods such as ZephIR [13] and CeNDeR [20]. This result demonstrates that ASCENT can circumvent the traditional trade-offs between accuracy and annotation effort. Furthermore, ASCENT substantially outperforms other annotation-free methods like the original NeRVE tracker (82.9%) [9] and fDNC (79.1%) [19], highlighting the power of ASCENT’s synergistic feature integration and contextual refinement.

**Table 1:**
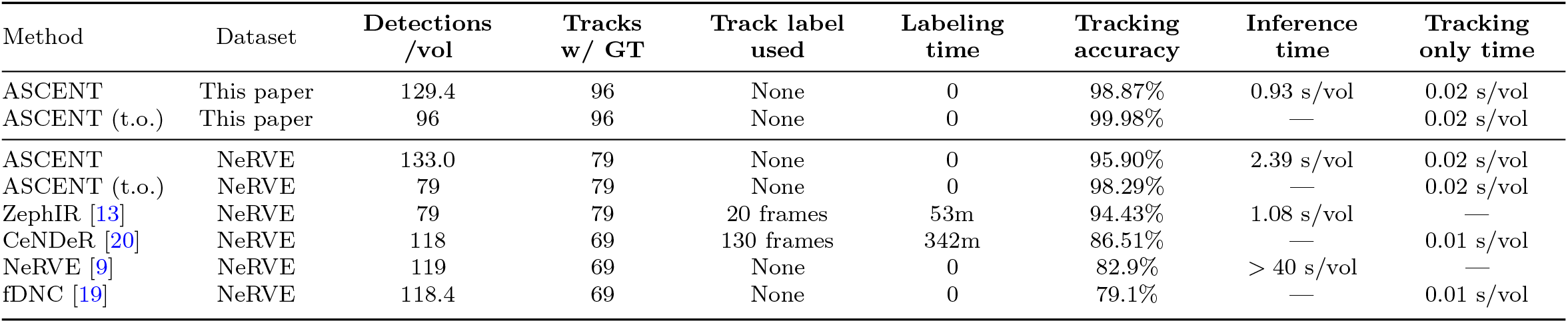
Benchmark evaluation of ASCENT compared to existing methods on the in-house and NeRVE datasets. “t.o.” (tracking only) refers to a processing mode where only ground-truth points are embedded and tracked. Labeling time is estimated based on the original NeRVE publication [9], which reported an annotation rate of approximately 2 seconds per neuron. Accuracy is measured using Modified MOTA (Equation 3).

| Method | Dataset | Detections<br>/vol | Tracks<br>w/ GT | Track label<br>used | Labeling<br>time | Tracking<br>accuracy | Inference<br>time | Tracking<br>only time |
| --- | --- | --- | --- | --- | --- | --- | --- | --- |
| ASCENT | This paper | 129.4 | 96 | None | 0 | 98.87% | 0.93 s/vol | 0.02 s/vol |
| ASCENT (t.o.) | This paper | 96 | 96 | None | 0 | 99.98% | — | 0.02 s/vol |
| ASCENT | NeRVE | 133.0 | 79 | None | 0 | 95.90% | 2.39 s/vol | 0.02 s/vol |
| ASCENT (t.o.) | NeRVE | 79 | 79 | None | 0 | 98.29% | — | 0.02 s/vol |
| ZephIR [13] | NeRVE | 79 | 79 | 20 frames | 53m | 94.43% | 1.08 s/vol | — |
| CeNDeR [20] | NeRVE | 118 | 69 | 130 frames | 342m | 86.51% | — | 0.01 s/vol |
| NeRVE [9] | NeRVE | 119 | 69 | None | 0 | 82.9% | > 40 s/vol | — |
| fDNC [19] | NeRVE | 118.4 | 69 | None | 0 | 79.1% | — | 0.01 s/vol |

**Fig. 4:**
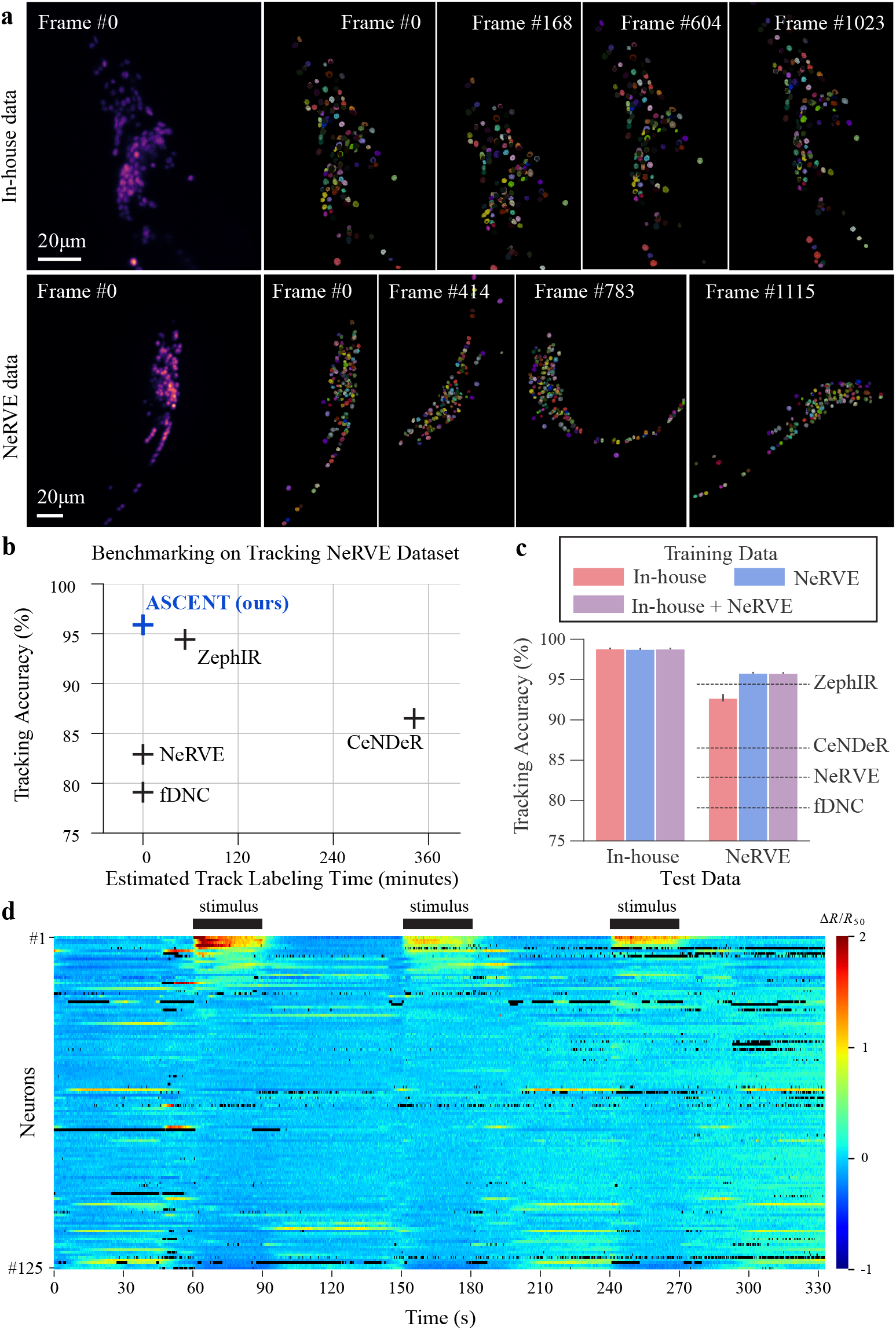
ASCENT achieves robust, SOTA performance and generalizes across diverse datasets. **a**, Example frames from the NeRVE and in-house datasets, illustrating differences in imaging conditions and animal confinement. **b**, Tracking accuracy versus manual labeling time required for various methods on the NeRVE benchmark. ASCENT achieves the highest accuracy with zero labeling time. Labeling time for other methods is estimated based on the original NeRVE publication [9], which reported an annotation rate of approximately 2 seconds per neuron. **c**, Cross-dataset generalization performance. Bars show the tracking accuracy of different training and test dataset combinations. Dotted lines indicate the performance of competitor methods on the NeRVE dataset for reference. **d**, Ratiometric neural activity traces (Δ*R/R*_50_) extracted from the in-house dataset using ASCENT’s tracking results. Attractive food odor stimuli were applied during 60–90s, 150–180s, and 240–270s.

A key hallmark of a robust computational method is its ability to generalize across different datasets and imaging conditions. We assessed the model’s ability to generalize by training it on one dataset and testing it on the other. A model trained on the more challenging, diverse NeRVE dataset generalized effectively to the in-house data, reaching an accuracy of 98.80%, which is nearly identical to the 98.86% achieved by a model trained specifically on the in-house data (Figure 4c). This strong crossdataset performance indicates that ASCENT learns fundamental, transferable features of neuron appearance and spatial configuration, rather than overfitting to the specific motion patterns of a single recording. Furthermore, a single model trained on a combination of both datasets performed robustly on each, achieving accuracy comparable to models trained on each specific dataset (Figure 4c). This is a non-trivial challenge, as naively training on mixed datasets can lead to performance trade-offs, where a model’s accuracy on a more difficult subset is compromised by the inclusion of simpler data [37, 38]. ASCENT, however, avoids this common pitfall. The combined model sustained high accuracy on both datasets, indicating that ASCENT learns a generalized feature space that transcends dataset-specific characteristics. This suggests the feasibility of creating a single, pre-trained ‘foundation model’ for neuron tracking that can be broadly applied to new datasets with minimal or no fine-tuning, greatly enhancing the method’s accessibility and utility.

The high-fidelity tracks produced by ASCENT enable reliable downstream analysis. For example, using the tracks from the in-house dataset, we could extract ratiometric neural activity traces that clearly reveal odor-entrained responses as well as spontaneous activities in the animal (Figure 4d; see Methods). To rigorously evaluate the fidelity of these tracks and pinpoint sources of error, we quantified the rates of missing and mismatched neurons for each frame. We found that both missing and mismatch errors are sparse, with most frames having zero or one error of each category. To isolate the source of error, whether from the core embedding and linking module, or from the upstream detection task, we evaluated ASCENT in a ‘track-only’ mode, where it processed ground-truth neuron coordinates directly (see Methods for details) (Table 1). The near-perfect accuracy in this mode (99.98%) compared to the standard mode (98.87%) on the in-house data confirms that the vast majority of errors in the end-to-end pipeline are attributable to the initial detection step. A qualitative analysis of the most error-prone frames supports this conclusion: errors often arise from segmentation failures, such as under-segmentation of neurons or ambiguities between ground-truth annotations and segmentation masks, rather than from the core embedding-and-linking methodology of ASCENT (Supplementary Figure 4). Improving the detection accuracy would further improve the end-to-end performance.

### High data efficiency enables rapid training on minimal data

Beyond its accuracy, a key practical advantage of ASCENT is its data efficiency. This is critical for applications such as rapidly training models for new experimental conditions or building large, foundational models from diverse recordings. To quantify this feature, we investigated the impact of training set composition on performance using the NeRVE dataset (Figure 5). A visualization of the image space across the entire recording reveals a circular and continuous manifold, suggesting considerable informational redundancy between temporally adjacent frames (Figure 5a; see Methods for visualization details). To test this, we created training sets of increasing size (1, 2, 4, …, 64 frames) by sampling frames at regular intervals across the recording. For comparison, we created an additional training set using only the first 8 consecutive frames (Figure 5b). We found that ASCENT achieves over 95% accuracy using as few as 8 regularly sampled, unlabeled frames, reaching near-peak performance with only a fraction of the full 1,532-frames dataset (Figure 5c). Notably, a model trained on the first 8 consecutive frames, which cover a less diverse range of postures, still achieved a high accuracy of 95.42%, only slightly lower than the 95.66% from 8 regularly sampled frames, suggesting that our strong data augmentation strategy can largely compensate for a lack of diversity in the training frames. This efficiency extends to training time, with convergence to plateau accuracy achieved in under 15 minutes with 1 H200 GPU for all tested subset sizes, highlighting the feasibility of quickly generating dataset-specific models without a significant time investment (Figure 5d). Although this data efficiency is advantageous, the self-supervised nature of ASCENT means that using all available frames to maximize exposure to diverse postures is always an option, as it incurs no additional manual labeling cost.

**Fig. 5:**
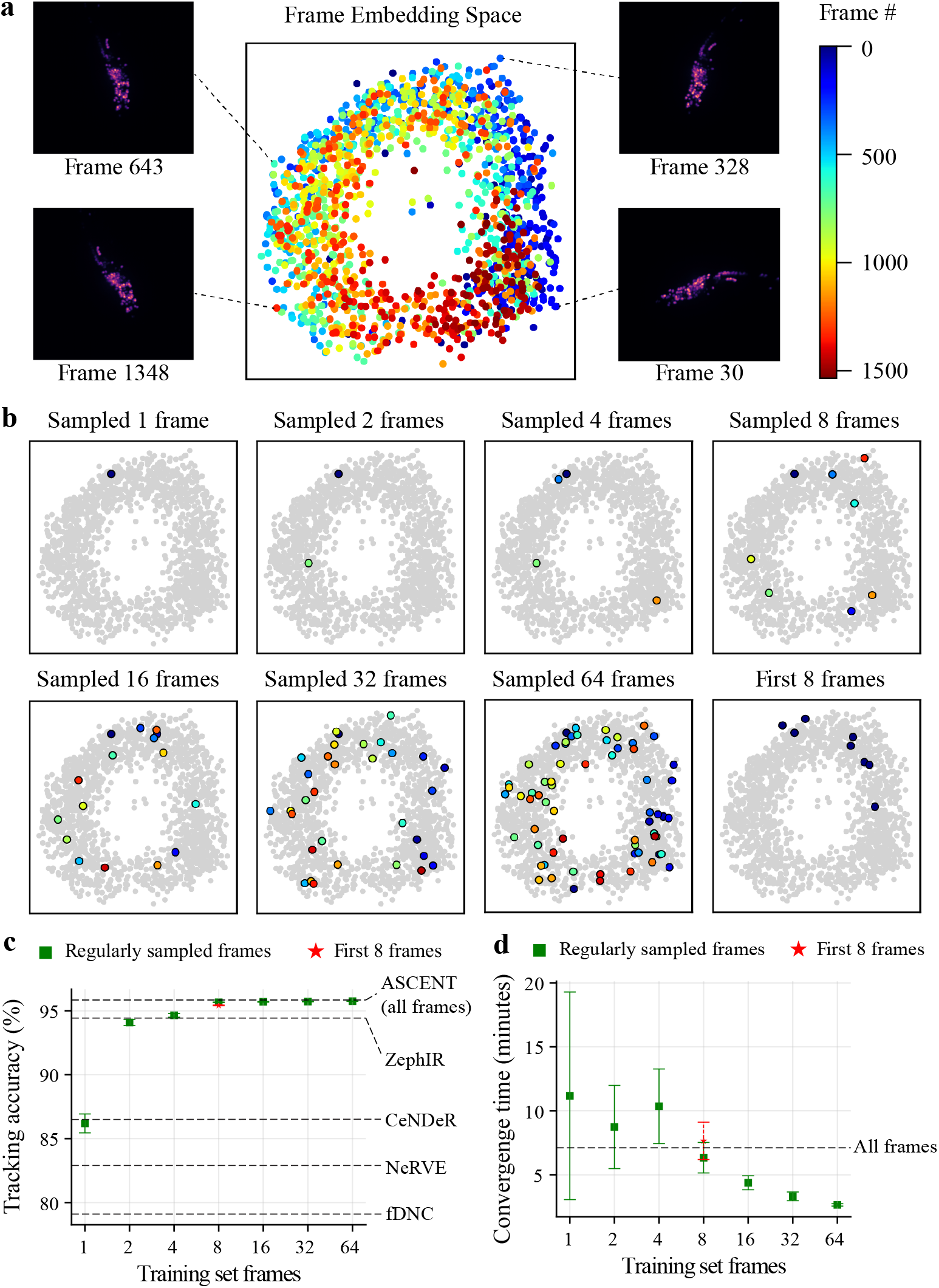
High data efficiency enables rapid model training on minimal data. **a**, t-SNE visualization of the image space of all frames from the NeRVE dataset. Each point represents a single frame, colored according to its point in time based on the color bar (right). Inset images show representative worm postures from different regions of the space. **b**, Visualization of the subsets of frames used for training. Sampled frames for each condition are highlighted in color against the full set of frames (gray). **c**, Tracking accuracy as a function of the number of training frames used. Accuracy plateaus quickly, achieving *>*95% with as few as 8–16 frames. The performance of competitor methods is shown as horizontal dashed lines for reference. **d**, Wall-clock convergence time as a function of training set size. Training is rapid across all conditions, typically requiring less than 15 minutes to reach convergence. Data are from the NeRVE dataset; error bars in **c** and **d** represent mean *±* standard deviation. over three independent runs.

## Discussion

In this work, we present ASCENT, a self-supervised framework that sets a new standard for annotation-free 3D neuron tracking. By training the Neuron Embedding Transformer, NETr, with a contrastive learning objective, ASCENT generates highly discriminative embeddings directly from unlabeled volumetric imaging data. This eliminates the most significant bottleneck in conventional tracking pipelines: the need for laborious manual track annotation. Our results demonstrate that the accuracy of this approach exceeds that of state-of-the-art methods that rely on supervised training.

A key insight from our architectural analysis is the effectiveness of learned local visual features. Contrary to the long-held assumption that the visual similarity of neurons necessitates a primary reliance on positional information, our results show that a model using only visual features can achieve remarkable accuracy on standard benchmark data, as long as noise is not too severe. This underscores the power of combining a strong pretrained visual backbone with a self-supervised objective tailored for instance discrimination. However, our experiments also confirm that this approach is not flawless; incorporating explicit positional encoding is critical for providing robustness against image noise often encountered in fluorescence imaging. Therefore, the integration of both visual and positional information within the NETr architecture offers the most reliable and broadly generalizable solution.

While optimizing neuron segmentation and detection models was beyond the scope of this work, ASCENT’s modularity means it can readily benefit from future advances in faster and more accurate keypoint detection or segmentation algorithms. Furthermore, while ASCENT’s track linking is highly efficient, scaling to datasets with tens of thousands of neurons, such as those from larger vertebrate brains, would require integrating more advanced, scalable assignment algorithms to overcome the computational complexity of exhaustive pairwise matching.

Looking forward, the principles demonstrated in ASCENT open several promising avenues for advancing the analysis of neural dynamics. The high data efficiency and strong generalization capabilities position ASCENT as a candidate for the development of a ‘foundation model’ for neuron tracking. Such a model could be pretrained on large, diverse benchmark datasets and then rapidly fine-tuned for specific experimental conditions, greatly increasing accessibility and standardization across the field. By removing the barrier of manual annotation and demonstrating high efficiency in both model training and inference, ASCENT makes high-performance 3D neuron tracking more accessible. This advance makes the quantitative analysis of entire neural systems more scalable, robust, and accessible, empowering researchers to tackle larger and more complex questions about the relationship between neural circuit activity and behavior.

## Methods

### Datasets and ground-truth

Experiments were conducted on two distinct *C. elegans* whole-brain imaging datasets. For all experiments, the stable red fluorescent channel, which tags all neuron nuclei, was used for neuron detection and for generating the embeddings used in tracking.

1. **NeRVE Dataset [3]:** This publicly available dataset features a freely moving animal (*C. elegans* strain AML14, expressing pan-neuronal GCaMP6s and tagRFP) and serves as a challenging benchmark due to significant inter-frame movements and non-rigid deformations. The dataset was acquired using a spinning disk confocal microscope equipped with a tracking system to keep the worm centered in the field of view [3]. It consists of 1,532 volumetric frames (2 × 34 × 600 × 632; channels × z × y × x) acquired at 6 Hz with a spatial resolution of 1.5*µm* in *z* and 0.3226*µm* in *xy*. Evaluation was performed against the manual track annotations provided by the original authors [3], which cover 79 neurons. This dataset is available at https://ieee-dataport.org/open-access/tracking-neurons-moving-anddeforming-brain-dataset.
2. **In-house Dataset:** This dataset was acquired to test performance under different imaging and behavioral conditions. It comprises a recording of neuronal activity across the head ganglion of *C. elegans* (strain ZM9624, expressing pan-neuronal GCaMP6 and mNeptune) confined within a microfluidic device based on the ‘olfactory chip’ design [39] to permit controlled chemical stimulation. Worms were subjected to repeated cycles of 60s buffer followed by a 30s OP50 supernatant stimulus. Imaging was performed on a Bruker Opterra II swept-field confocal microscope using a 40x, 0.75 NA air objective and a Photometrics Evolve 512 Delta EM-CCD camera. The recording consists of 1,100 volumetric frames (2 × 9 × 512 × 512; channels × z × y × x) acquired at 3.3 Hz with a spatial resolution of 1.5*µm* in *z* and 0.243*µm* in *xy*. Ground-truth tracks for 96 neurons were generated by manually proofreading and correcting an initial set of tracks generated by ZephIR [13].

### Data pre-processing

A key advantage of ASCENT is its minimal pre-processing pipeline. Unlike methods that require computationally expensive image registration, animal straightening, or graph construction, ASCENT operates directly on normalized volumetric data. We apply percentile normalization on a frame-by-frame basis, where intensity values are linearly rescaled such that the 1st and 99.99th percentiles of each frame’s voxel intensities map to a range of [0, 1]. This approach effectively mitigates the impact of outlier pixels (e.g., dead or hot pixels) without distorting the intensity distribution of the relevant neuronal signals.

### Problem formulation

ASCENT employs a tracking-by-detection strategy. Given a volumetric time-series recording, *I* ∈ ℝ^*T* ×*C*×*D*×*H*×*W*^, where *T* is the number of frames, *C* the number of channels, and *D, H*, and *W* are the spatial dimensions, the framework proceeds as follows. For each frame *I*_*t*_, a detection algorithm provides a set of *n*_*t*_ neuron candidates 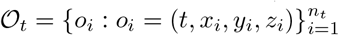, where (*x*_*i*_, *y*_*i*_, *z*_*i*_) are the spatial coordinates. The core of the framework is the Neuron Embedding Transformer (NETr), denoted as *f*_*θ*_(·), which maps the image data *I*_*t*_ and the set of detections 𝒪_*t*_ to a set of discriminative feature vectors 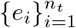, where each embedding *e*_*i*_ ∈ ℝ^*d*^:

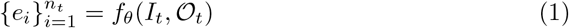

These embeddings *e*_*i*_ serve as the basis for linking detections across time frames to build a set of tracks, *TR* = {*tr*_1_, *tr*_2_, …, *tr*_*K*_}, where *K* is the total number of unique neuron trajectories. Each track *tr*_*k*_ represents the trajectory of a single neuron, defined as a sequence that associates a persistent identity *k* with at most one neuron detection from *O*_*t*_ for each frame *t* ∈ {1, …, *T*}.

### Neuron detection

Neuron candidate coordinates for each frame were obtained using a custom-trained 3D StarDist [40] model specific to each dataset. To generate the necessary training data, ground-truth instance masks were created for two representative frames from each dataset. This involved first applying a pre-trained nnUNet [41] model to produce an initial binary segmentation mask that distinguishes neuron pixels from the background. Connected component analysis was then performed on this output to generate preliminary instance labels. These labels were subsequently manually proofread and corrected to ensure accurate, individual neuron identities suitable for training. Each dataset-specific StarDist model was then trained for 500 epochs using these curated instance masks. The 3D coordinates of the final detected neuron centroids served as the positional input, {*o*_*i*_}, to the NETr embedding network.

### NETr architecture

The Neuron Embedding Transformer (NETr) is designed to generate context-aware embeddings by combining local visual appearance and positional information, refined through a self-attention mechanism. The network processes batches of image frames and their corresponding neuron detections to produce a set of unique embeddings.

The process begins with local visual feature extraction. For each detected neuron *o*_*i*_ in a batch of frames, a 3D local patch of size (*p*_*x*_, *p*_*y*_, *p*_*z*_) = (64, 64, 3) pixels is extracted around its centroid coordinates. To achieve high efficiency, this operation is vectorized across all neurons in the batch using a pre-computed sampling grid. The three z-slices of each patch are treated as the input channels for a 2D ChannelViT backbone, a Vision Transformer variant adapted for multi-channel images [42]. This backbone, pre-trained on the ImageNet-1k dataset using the DINO self-supervised learning method [35], outputs a feature vector, *vis*_*i*_, which is then projected to the target embedding dimension, *d* = 256. Concurrently, positional features are generated. The 3D coordinates of each neuron are normalized and processed by a 3-layer multilayer perceptron (MLP), which is implemented as a 1D convolutional network for computational efficiency, with Batch Normalization applied between layers. This module produces a positional embedding, *pos*_*i*_ ∈ ℝ^*d*^. These two feature vectors are then combined via element-wise addition to yield an initial representation for each neuron: *h*_*i*_ = *vis*_*i*_ + *pos*_*i*_. Finally, the sequence of initial representations, 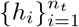, for each frame undergoes contextual refinement using a 4-layer Transformer Encoder [43]. The encoder employs self-attention with 4 attention heads, a feedforward dimension of 512, and a dropout rate of 0.1. A key padding mask is generated from the input neuron identifiers to ensure that padded, non-existent neurons are ignored during the self-attention calculation. The output from the encoder provides the final, contextually refined embeddings {*e*_*i*_}, with a final embedding dimension *d* of 256.

### Self-supervised contrastive training of NETr

To learn discriminative embeddings without manual track annotations, we employ a self-supervised contrastive learning strategy based on the SimCLR framework [31]. The objective is to train the NETr network, *f*_*θ*_, to produce embeddings that are invariant to data augmentations while remaining specific to each neuron’s identity.

The training proceeds in mini-batches, where each batch contains *B* frames randomly selected from the dataset. For each frame *t* in the batch, consisting of the image *I*_*t*_ and its *n*_*t*_ detections *O*_*t*_, we apply two independent random augmentation pipelines, *ϕ*_1_ and *ϕ*_2_. This process generates two distinct augmented “views” of the same set of neurons, (*I*_*t*,1_, *O*_*t*,1_) and (*I*_*t*,2_, *O*_*t*,2_). These augmentations include transformations relevant to 3D microscopy, such as rotation, elastic deformation, resized cropping, position jitter, intensity variations, and object dropout. Both augmented views for the entire batch are passed through the NETr network to produce two sets of embeddings, ℰ^(1)^ and ℰ^(2)^. The network parameters *θ* are then optimized by minimizing the NT-Xent (Normalized Temperature-scaled Cross-Entropy) loss [31], which is computed for each frame in the batch and then averaged. For a single frame with *N*^*′*^ valid neurons present in both augmented views, the loss for a positive pair of embeddings (*e*_*i*_, *e*_*j*_), corresponding to the same neuron *i* under two different augmentations, is defined as:

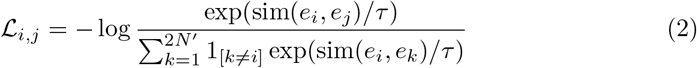

where *sim*(*u, v*) is the cosine similarity between vectors *u* and *v, τ* is a temperature hyperparameter set to 0.05, and the indicator function 1_[*k*≠*i*]_ ensures the denominator sums over all embeddings in the augmented set of 2*N*^*′*^ embeddings except for the anchor *e*_*i*_ itself. This objective function attracts embeddings of the same neuron (positive pairs) while repelling embeddings of different neurons (negative pairs). The final loss for the frame is averaged over all positive pairs, computed symmetrically for both views.

In this study, models were trained using the Adam optimizer with an initial learning rate of 1 × 10^−4^ and a batch size of 4 frames. All training was performed on a single NVIDIA H100 GPU. The number of training epochs varied depending on the experiment: 20 epochs were used for the benchmark and generalization experiments, 300 epochs for the augmentation ablation study, and the epoch count was adjusted for the data-efficiency study to normalize the total number of training steps.

### Track-linking algorithm

The discriminative embeddings produced by NETr enable a simplified track-linking stage that does not require complex post-processing heuristics. After generating embeddings {*e*_*i*_}_*t*_ for all detections in each frame *t*, tracks were formed using serial bipartite matching. For each subsequent frame, a cost matrix 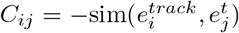, where 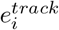 is the current embedding of an active track and sim is the cosine similarity, was computed between active tracks and new detections. A dynamic cost threshold *θ*_*t*_ was estimated as 0.5 × (mean(top-*k* costs) +mean(all costs)), where *k* is the number of active tracks. The Hungarian algorithm was used to find the minimum cost assignment, and any potential matches with a cost *C*_*ij*_ *> θ*_*t*_ were rejected. Detections that remained unmatched initiated new tracks. The embeddings of successfully matched tracks were updated using an exponential moving average: 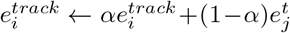, with a momentum parameter *α* = 0.5.

### Track-only mode

To isolate the performance of the core embedding and linking module from errors introduced by the upstream neuron detection step, we evaluated ASCENT in a ‘track-only’ mode. In this mode, instead of using the output of a detection algorithm, the ground-truth neuron coordinates from the manually annotated track labels were used as the input detections, 𝒪_*t*_, for the NETr embedding network. This approach provides a direct measure of the embedding and linking performance by removing the confounding variable of detection inaccuracies, such as missed neurons or imprecise localizations. The subsequent tracking and evaluation procedures remained identical to the standard pipeline.

### Ablation studies

1. **Architecture ablation:** To dissect the NETr architecture, we created two ablated models: a ‘Visual features only’ model that omitted the positional encoding MLP, and a ‘Position features only’ model that omitted the ChannelViT visual backbone. These were compared against the ‘Standard’ full model. To test for robustness, models were trained and evaluated on modified versions of the NeRVE dataset where Gaussian noise with a mean of 0 and a standard deviation (*σ*) of 0.1, 0.2, and 0.3 was added to the normalized image data.
2. **Augmentation ablation:** To evaluate the contribution of different data augmentations, the standard NETr model was trained on a reduced dataset of 16 regularly sampled NeRVE frames for 300 epochs. Performance was systematically tested by either including only one or two augmentations at a time, or by starting with the full suite of six augmentations and removing one or two at a time.

### Tracking-evaluation metrics

Tracking accuracy was evaluated following the methodology of previous works [9, 13, 20]. Correspondences between ground-truth points and pipeline-tracked points were established by finding mutually closest pairs within a maximum Euclidean distance of 5*µm*. Based on these matches, we defined two primary error types: Missing errors, where a ground-truth point has no matched point in the tracker output, and Mismatch errors (ID switches), where a tracked point is assigned a track ID that differs from its matched ground-truth point. We use a modified Multiple Object Tracking Accuracy (MOTA [44]) metric that excludes false positives, as ground-truth annotations typically cover only a known subset of all neurons present:

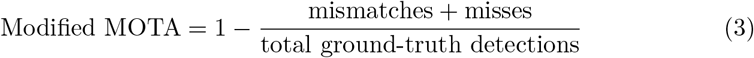

This metric reflects the tracker’s ability to correctly maintain neuron identities over time relative to the provided ground-truth.

### UMAP visualization of the neuron embedding space

To visualize the structure of the learned feature space, we used the uniform manifold approximation and projection (UMAP) algorithm [36]. We fitted two separate 2D UMAP models to project the high-dimensional neuron embeddings into a 2D space. The first model was fitted using embeddings generated by the randomly initialized NETr network (before training), and the second was fitted using embeddings from the fully trained network. To train each UMAP model, we first generated feature embeddings for all ground-truth neurons across all frames of the NeRVE dataset using the corresponding (untrained or trained) NETr network. Cosine similarity was specified as the distance metric for the UMAP algorithm, consistent with the similarity metric used in the contrastive training objective. For the visualizations shown in the main text, embeddings from an example frame and its randomly augmented counterpart were then projected into these pre-fitted 2D UMAP spaces.

### t-SNE visualization of the frame space

To visualize the global image space throughout the NeRVE recording, we projected each frame into a low-dimensional space using a two-stage process. First, we trained a convolutional autoencoder to learn a compressed 32-dimensional representation for each frame. The input data for this network was generated by computing a maximum intensity projection along the z-axis for each of the 1,532 volumetric frames, with each resulting 2D image resized to 64 × 64 pixels. The autoencoder, consisting of a convolutional encoder and a corresponding decoder, was trained for 1,000 epochs using the Adam optimizer (learning rate = 1 × 10^−3^) to minimize the mean squared error (MSE) between the input and reconstructed images. Next, the trained encoder was used to extract a 32-dimensional latent vector for every frame in the dataset. We then applied t-Distributed Stochastic Neighbor Embedding (t-SNE) to these latent vectors to generate a 2D embedding suitable for visualization.

### Extraction of calcium activity

To generate ratiometric neural activity traces, we used the two-channel fluorescence data from the GCaMP (green) and structural marker (red) channels. For each tracked neuron at each time point, fluorescence intensity was calculated independently for each channel. A 7 × 7 × 3 pixel (x, y, z) patch was defined around the neuron’s centroid, and the raw fluorescence signal (*F*_*G*_ for green, *F*_*R*_ for red) was determined by averaging the values of the 10 brightest pixels within this patch. A channel-specific background value, calculated as the modal intensity of each frame, was then subtracted from the raw signal. The background-corrected intensities were used to compute a ratiometric signal, 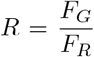, for each time point. For each neuron, a baseline fluorescence ratio, *R*_50_, was established by taking the median of its ratiometric signal *R* over the full duration of the video recording. Finally, 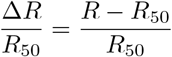 was calculated and plotted.

## Data availability

The datasets generated and/or analyzed during the current study will be available upon peer-reviewed publication. The NeRVE dataset is publicly available as cited.

## Code availability

The codes implemented for this study will be available upon peer-reviewed publication.

## Supplementary information

Supplementary Figure 1, 2, 3, 4.

---

**Supplementary Fig. 1:**
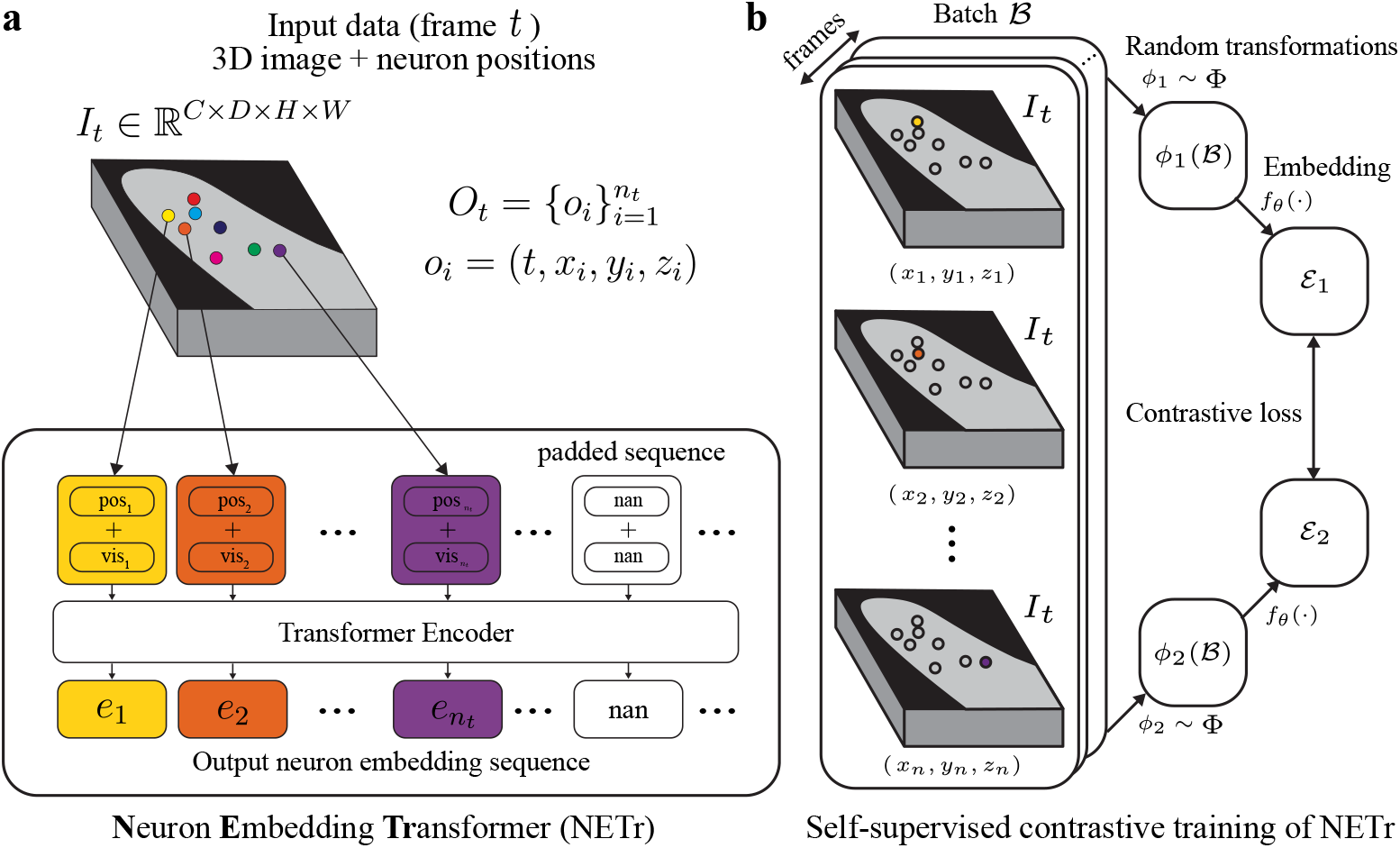
Detailed architecture and training scheme of the Neuron Embedding Transformer (NETr). **a**, The NETr architecture. For each detected neuron, NETr combines local visual features (*vis*_*i*_) with positional features (*pos*_*i*_). Visual features are extracted from a local 3D patch using a DINO-pretrained ChannelViT backbone. Positional features are generated from the neuron’s 3D coordinates using a multi-layer perceptron (MLP). A Transformer Encoder then uses self-attention to integrate contextual information from all neurons in the frame 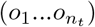 to produce final, refined embeddings (*e*_*i*_). **e**, The self-supervised training scheme. The network is trained by applying two independent random transformations (*ϕ*_1_, *ϕ*_2_) to batches of frames. A contrastive loss function maximizes the similarity between embeddings of the same neuron 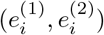 from the two augmented views, while minimizing similarity between different neurons.

**Supplementary Fig. 2:**
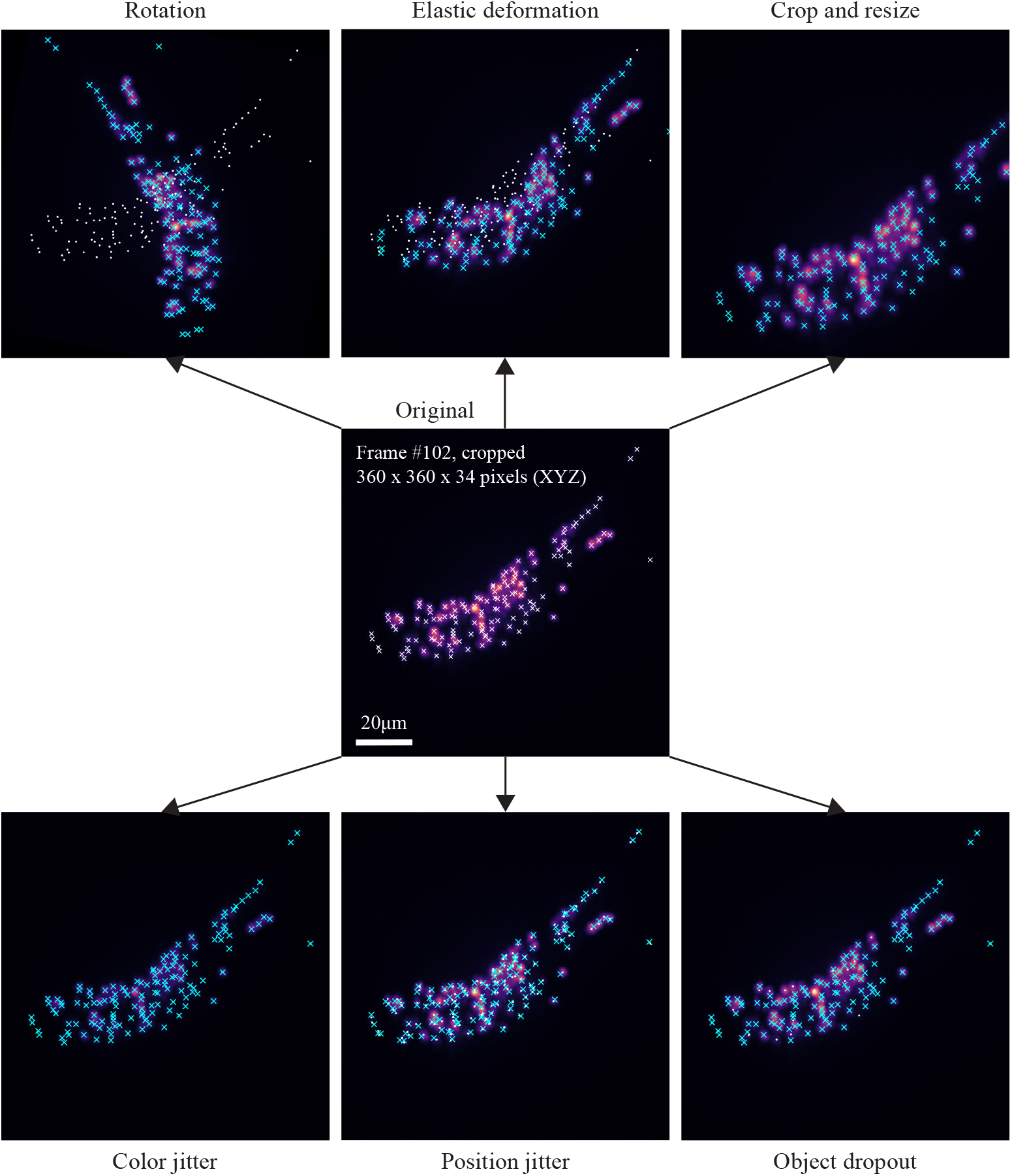
Data augmentations for self-supervised learning of NETr. Examples of the six individual transformations applied to an original frame from the NeRVE dataset (center), shown as maximum intensity projections of a 360 × 360 pixel cropped region. In the original image, detected neurons are marked with white ‘ ×’s. In the transformed images, the new neuron positions are marked with cyan ‘ ×’s, while the original positions are shown with white dots where applicable to illustrate the transformation’s effect. Geometric transformations such as rotation, elastic deformation, and crop/resize alter both image and coordinate data. Color jitter modifies only the image data, while position jitter and object dropout alter only the coordinate data.

**Supplementary Fig. 3:**
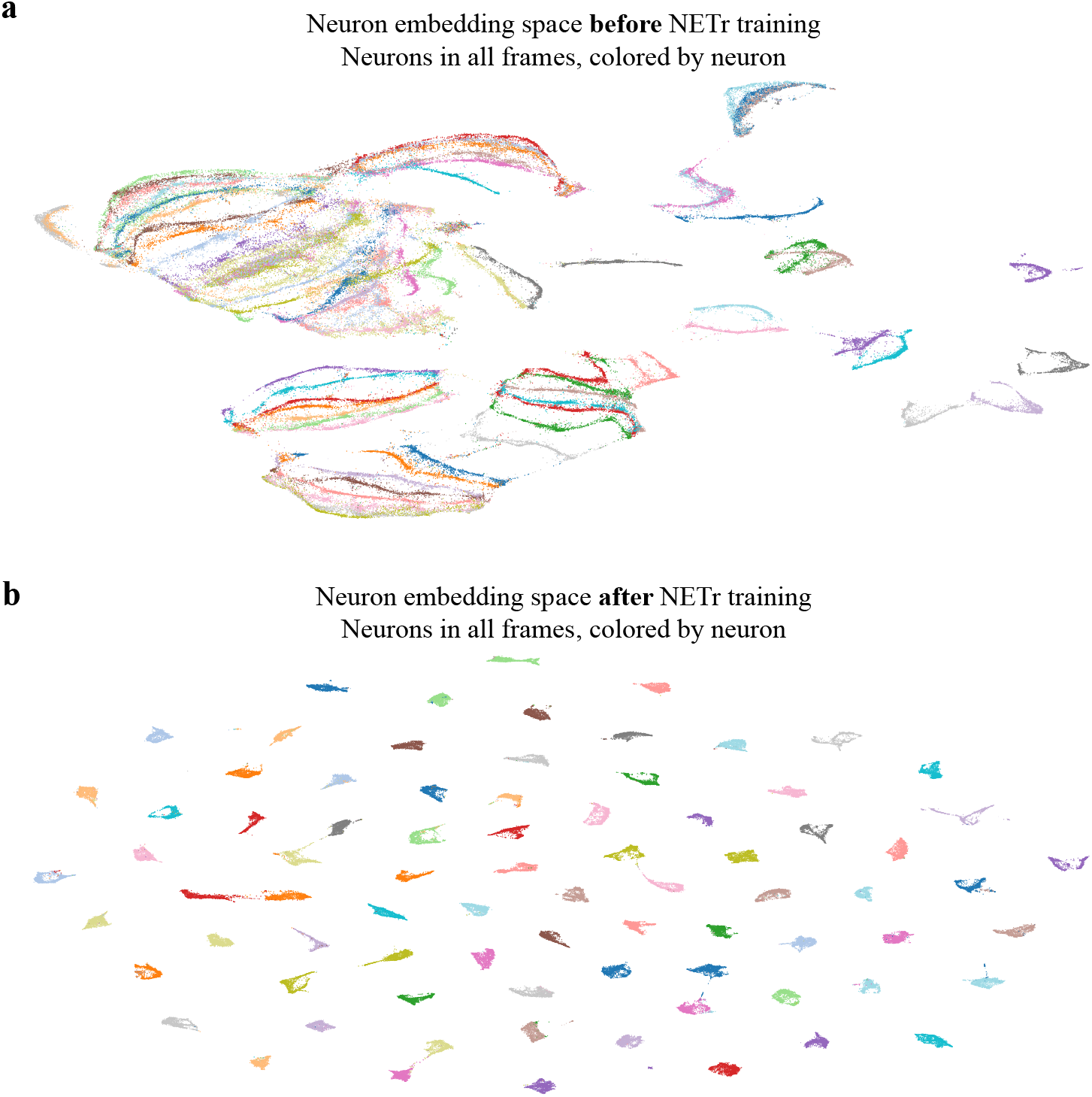
UMAP visualization of the neuron embedding space. High-dimensional feature embeddings for all ground-truth neurons across all frames of the NeRVE dataset were projected into a 2D space using UMAP. Each point represents a single neuron at a single timepoint, and points are colored according to their unique neuron identity. **a**, Before training, the embeddings are unstructured and intermingled, indicating the network cannot distinguish between different neurons. **b**, After self-supervised contrastive training, the embeddings form tight, identity-specific clusters, demonstrating that the network has learned a discriminative feature space where each neuron identity occupies a unique region of the space.

**Supplementary Fig. 4:**
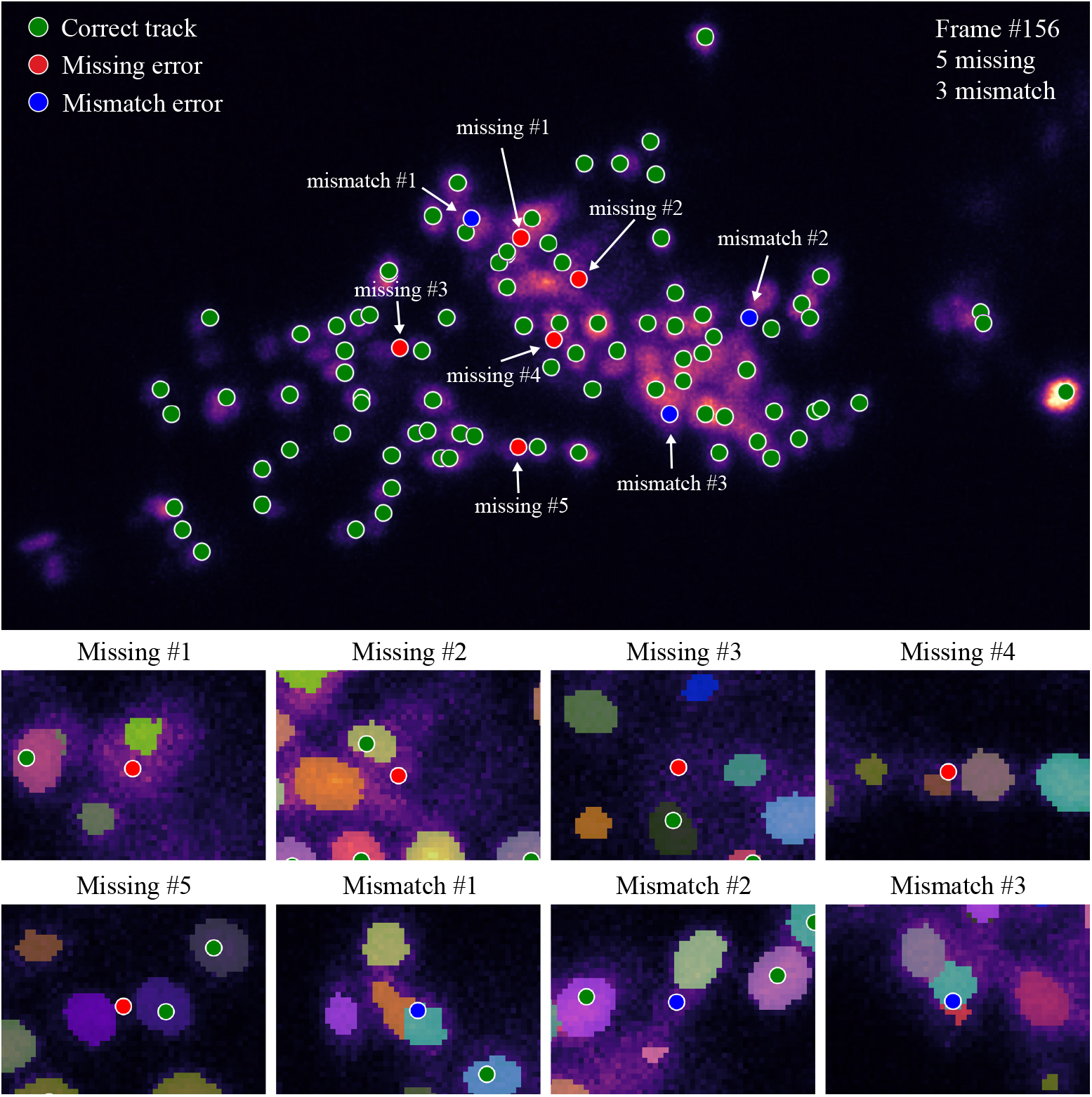
Qualitative analysis of tracking errors. Detailed breakdown of tracking errors in an error-prone frame from the in-house dataset (frame #156). The main panel shows a maximum intensity projection of the frame, with ground-truth neurons colored by their tracking status: correctly tracked (green), missing (red), or mismatched (blue). The eight bottom panels show zoomed-in views of individual errors at the relevant z-plane, with the StarDist-generated segmentation mask overlaid. The analysis reveals that the majority of errors are attributable to the upstream detection step. Missing #2, #3 are due to StarDist failing to detect the neurons. Missing #1, #4, #5 and mismatch #2 are caused by incomplete segmentation masks that do not sufficiently cover the ground-truth neuron position. Mismatch #1, #3 occur where StarDist detects two distinct objects near a single ground-truth neuron, suggesting potential ambiguities in the ground-truth annotation itself.

